# Exploring valence bias as a metric for frontoamygdalar connectivity and depressive symptoms in childhood

**DOI:** 10.1101/839761

**Authors:** Nathan M. Petro, Nim Tottenham, Maital Neta

## Abstract

Negativity bias is a core feature of depression that is associated with dysfunctional frontoamygdalar connectivity; this pathway is associated with emotion regulation and sensitive to neurobiological change during puberty. We used a valence bias task (ratings of emotional ambiguity) as a potential early indicator of depression risk and differences in frontoamygdalar connectivity. Previous work using this task demonstrated that children normatively have a negative bias that attenuates with maturation. Here, we test the hypothesis that persistence of this negativity bias as manturation ensues may reveal differences in emotion regulation development, and may be associated with increased risk for depression. Within a restricted age range (6-13 years), we tested the moderating role of puberty on relationships between valence bias, depressive symptoms, and frontoamygdalar connectivity. A negative bias was associated with increased depressive symptoms for those at more advanced pubertal stages (within this sample) and less regulatory frontoamygdalar connectivity, whereas a more positive bias was associated with more regulatory connectivity patterns. These data suggest that with maturation, individual differences in positivity biases and associated emotion regulation circuitry confer a differential risk for depression. Longitudinal work is necessary to determine the directionality of these effects and explore the influence of early life events.

## 1. Introduction

Individual differences in internalizing problems, like depression, are associated with amygdala and medial prefrontal cortex (mPFC) circuitry (Gee, Gabard-Durnam, et al., 2013; Pezawas et al., 2005; Wang et al., 2009), and have been shown to emerge during the transition from childhood to adulthood (Emslie et al., 2005; Hankin et al., 1998; Kessler et al., 2001). Notably, a number of studies have linked the emergence of these individual differences to neurobiological changes during pubertal development (Andersen & Teicher, 2008; Angold & Costello, 2006; Paus et al., 2008). Indeed, developmental changes in the structure and function of amygdala-mPFC circuitry are intimately tied to the release of gonadal hormones during puberty (Lebron-Milad & Milad, 2012; van Wingen et al., 2011), and accompany profound changes in emotional behavior during this period of development that support functional social behavior (Denham, 1998; Eisenberg et al., 1993; John & Gross, 2004; Saarni, 1984).

In adulthood, connections between amygdala and frontal cortex are understood to underpin emotion regulation mechanisms (Banks et al., 2007; Phillips et al., 2008), where frontal regions downregulate amygdala activity during emotional processing (Hare et al., 2008; Hariri et al., 2003; Pezawas et al., 2005). Further, adults compared to children show relatively decreased amygdala activation (Guyer et al., 2008; Hare et al., 2008; Monk et al., 2003; Swartz et al., 2014; but see Moore et al., 2012) and greater inverse (i.e., regulatory) amygdala-mPFC connectivity (Gabard-Durnam et al., 2014; Gee, Humphreys, et al., 2013; Perlman & Pelphrey, 2011) in response to negative emotional information. Indeed, younger children show a non-regulatory positive amygdala-mPFC connectivity pattern, but older children and adolescents show an inverse regulatory connectivity suggestive of emotion regulation (Gee, Humphreys, et al., 2013). Moreover in adults, this more positive amygdala-mPFC connectivity has been implicated in the development of mental health disorders (Das et al., 2007; Gee, Gabard-Durnam, et al., 2013; Phillips et al., 2008).

One notable feature of internalizing symptoms is a negativity bias, or an enhanced attention to and memory for negative information (Browning et al., 2010; Roy et al., 2008; Vasey et al., 1995), which is central to a variety of mood and anxiety disorders (e.g., depression; Beck, 1976; Williams et al., 2007). Interestingly, negativity bias is associated with more positive amygdala-mPFC connectivity (i.e., a non-regulatory pattern) (Etkin et al., 2009; Roy et al., 2013; see also Kim et al., 2011). Given that this same circuitry is sensitive to pubertal changes (Blakemore et al., 2010; Goddings et al., 2014; Herting et al., 2014; Mills et al., 2014; Vijayakumar et al., 2018, 2019) and is related to mental health disorders which emerge during puberty (Burghy et al., 2012; Hulvershorn et al., 2011; Kessler et al., 2005; Lee et al., 2014; Roy et al., 2013), understanding how this circuitry is associated with individual differences in negativity bias across puberty is essential to understanding the development of depression.

Because the release of gonadal hormones which drive neurobiological changes can begin as early as 8 years (Blakemore et al., 2010; Dorn et al., 2006), whereas depression tends to onset at around 14 years of age (Burke, 1991; Lewinsohn et al., 1994), the construction of this system may begin in relatively early pubertal stages prior to the onset of internalizing symptoms. That is, pubertal transitions may act as a developmental “prism”, revealing individual differences in emotion regulation behaviors. In particular, the period from early to middle puberty is met with structural (Blanton et al., 2012; Goddings et al., 2014; Hu et al., 2013; Pfefferbaum et al., 2016) and functional (Clark & Beck, 2010; Forbes et al., 2011; Moore et al., 2012; Pfeifer et al., 2013; Vijayakumar et al., 2018, 2019) changes within both the amygdala and mPFC, as well as in their connectivity (Asato et al., 2010; Herting et al., 2014; Menzies et al., 2015; see also Gee et al., 2013). This evidence is consistent with detailed animal models which find that the onset of puberty triggers reorganization within both the mPFC and its white matter fibers shared with the amygdala (Juraska & Willing, 2017; Zimmermann et al., 2019). The neurobiological processes associated with these relatively early pubertal stages may produce measureable individual differences in frontoamygdalar function critical to mental health outcomes.

Reliably measuring emotional biases in children during these early pubertal stages poses a methodological challege. For example, extant literature on negativity bias, which includes subclinical symptomology (Pagliaccio et al., 2014), tends to measure emotional biases using stimuli that rely on developmentally advanced cognitive/linguistic abilities not ideally suited for younger children. An emerging alternative approach has measured emotional biases using valence ratings of facial expressions, which are reliably identified in early childhood (Bruce et al., 2000; Widen & Russell, 2008). Although some expressions (e.g., happy or angry) signal clear valence information about the emotions and intentions of others, other expressions (e.g., surprise) are ambiguous because they signal both positive (e.g., an unexpected gift) and negative events (e.g., witnessing an accident). Notably, children compared to adults tend to rate surprised expressions as having a more negative meaning (i.e., more negative valence bias; Tottenham, Phuong, Flannery, Gabard-Durnam, & Goff, 2013). Given that surprise expressions may be reliably identified across all ages and their ratings track developmental changes in emotional behavior, the ambiguity conveyed through surprised expressions is ideally suited to probe differences in negativity bias at early pubertal stages.

Neuroimaging work suggests that normative developmental shifts in emotional biases might reveal how pubertal changes correlate with variability in the neurobiology of depression. For instance, in adults, positive ratings of surprised faces depend upon slower and more deliberate processing (Kaffenberger et al., 2010; Neta et al., 2011; Neta & Tong, 2016), and appear to rely on emotion regulation mechanisms (Kim, Somerville, Johnstone, et al., 2003; Petro et al., 2018). This work suggests that the development of a more positive valence bias and a more mature (inverse) amygdala-mPFC connectivity during puberty can powerfully impact the emergence of internalizing symptoms during this developmentally sensitive period. For example, maintaining a negative valence bias and a more immature (positive) amygdala-mPFC connectivity into adulthood may be a risk factor for the onset of depression. However, no study has yet used neuroimaging to explore valence bias in youth populations. Thus, measuring normative shifts in valence bias and its associated amygdala-mPFC connectivity is an innovative approach that could reveal how pubertal changes correlate with variability in the neurobiology of depression.

Using a cross-sectional design, we explored the relationships between valence bias, depressive symptoms, and emotion regulation circuitry within an age range demonstrated to show a more negative valence bias (ages 6-13 years; Tottenham et al., 2013), a developing regulatory circuitry (Gabard-Durnam et al., 2014; Gee et al., 2013; Perlman & Pelphrey, 2011; Silvers et al., 2017), and prior to the typical emergence of depression (age 14 years; Burke, 1991; Lewinsohn et al., 1994). This methodological choice enabled us to explore how relatively early pubertal changes contribute to the emergence of internalizing problems in depression. We examined functional brain connectivity while viewing facial expressions during magnetic resonance imaging (MRI). We used pubertal scores as our proxy for maturity because (a) puberty is more closely tied than age to neurobiological changes in brain structure and function (Blakemore et al., 2010; Goddings et al., 2012) and especially amygdala-mPFC circuitry (Gabard-Durnam et al., 2014; Gee et al., 2013; Herting et al., 2014; Perlman & Pelphrey, 2011; Silvers et al., 2017), and (b) age represents a different stage in maturation for males and females (Schuiling & Likis, 2016). For our exploratory analysis, we predict that with increasing maturation (within this relatively immature sample), a more negative valence bias and a more immature amygdala-mPFC connectivity pattern may pose an increased risk for depression.

## 2. Materials and Methods

### 2.1 Participants

We collected data from 61 participants (29 female; ages 6-13 years, mean(SD) age = 9.18(2.13)). All participants and their parent reported that the participant had no history of neurological or psychiatric disorders, nor were any taking psychotropic medications. All protocols were approved by the University of Nebraska Committee for the Protection of Human Subjects. The participants and their parent were informed of all procedures prior to the child’s participation, and a parent of each participant gave written informed consent prior to testing in accordance with the Declaration of Helsinki.

Six participants did not complete the neuroimaging portion of the task. An additional fourteen participants were excluded for failing to accurately rate the clearly valenced angry (N=4) and happy (N=10) expressions on at least 60% of trials, an accuracy threshold used for adult participants (Brown et al., 2017; Neta et al., 2009, 2013, 2018; Neta & Tong, 2016). The exclusion criteria for accuracy is particularly important because, if participants are not accurately rating angry faces as negative and happy faces as positive, then it is difficult to discern the specific valence interpretations of emotional ambiguity (i.e., valence bias). The final sample consisted of 41 participants (23 female; ages 6-13 years, mean(SD) age = 9.46(2.12)) that did not differ in age from those excluded (*t_59_* = 1.44; *p* = .16; range = 6-13; mean(SD) age = 8.60(2.06)), and who identified as either Caucasian (N = 38), Black/African-American (N = 1), Asian (N = 1), or Unknown (N = 1).

Albeit relatively small given the analysis of individual differences (Dubois & Adolphs, 2016), this sample size allowed us to explore the potential relationship between pubertal development, depressive symptoms, valence bias, and amygdala-mPFC functional connectivity. Further, given that no study has combined this valence bias task with neuroimaging in childhood, this exploration aims to serve as a first step in developing a testable model of the development of depressive symptoms and emotion regulation circuitry as measured by the valence bias task.

#### 2.1.1 Individual difference measures

We assessed pubertal status using the scores from the Petersen Puberty Development Scale (PDS; Petersen, Crockett, Richards, & Boxer, 1988), which were calculated, as described by Petersen and colleagues, as the average of each of the 5 items (each with a possible response of 1-4) related to pubertal change (possible range = 1-4). Because PDS scores tend to deviate from 1 only after the age of 8 (Hu et al., 2013), all participants under the age of 8 were assigned the lowest score of 1 (i.e. prior to the onset of puberty). Because PDS score (Shapiro-Wilk: *W* = .90, *p* = .01), age (Shapiro-Wilk: *W* = .94, *p* = .03), and depressive symptoms (see below; Shapiro-Wilk: *W* = .78, *p* = .00001) were non-normally distributed, robust regression was used for all analyses of these variables.

As expected, given our targeted age range, the current sample subtended to relatively low PDS scores (range = 1-2.8, mean(SD) = 1.67(.56), skewness(SD) = .71(.37), kurtosis(SD) = - 0.54(0.72)) rather than representing the full spectrum of possible scores. This questionnaire was completed by the child with instruction from the experimenter and is comprised of questions related to secondary sex characteristics (e.g., growth of body hair, deepening of voice), the appearance of which are a marker of gonadal hormone release during gonadarche (Susman & Rogol, 2013). The PDS score was not related to sex (Figure 1A; *t*_39_ = −0.16, *p* = .87), but was positively corelated with age (Figure 1B; *t*_39_ = 6.68,*p* < .001). The average full-scale intelligence quotient of the sample was within normal range as assessed by the 2 subset Wechsler Abbreviated Scale of Intelligence (WASI; range = 88-142; mean(SD) = 114.18(14.53), however there were missing data for one participant; Wechsler, 1999), and was not related to age (*t*_38_ = - 0.92, *p* = .36) or PDS score (*t*_38_ = −0.66, *p* = .52).

**Figure 1.**
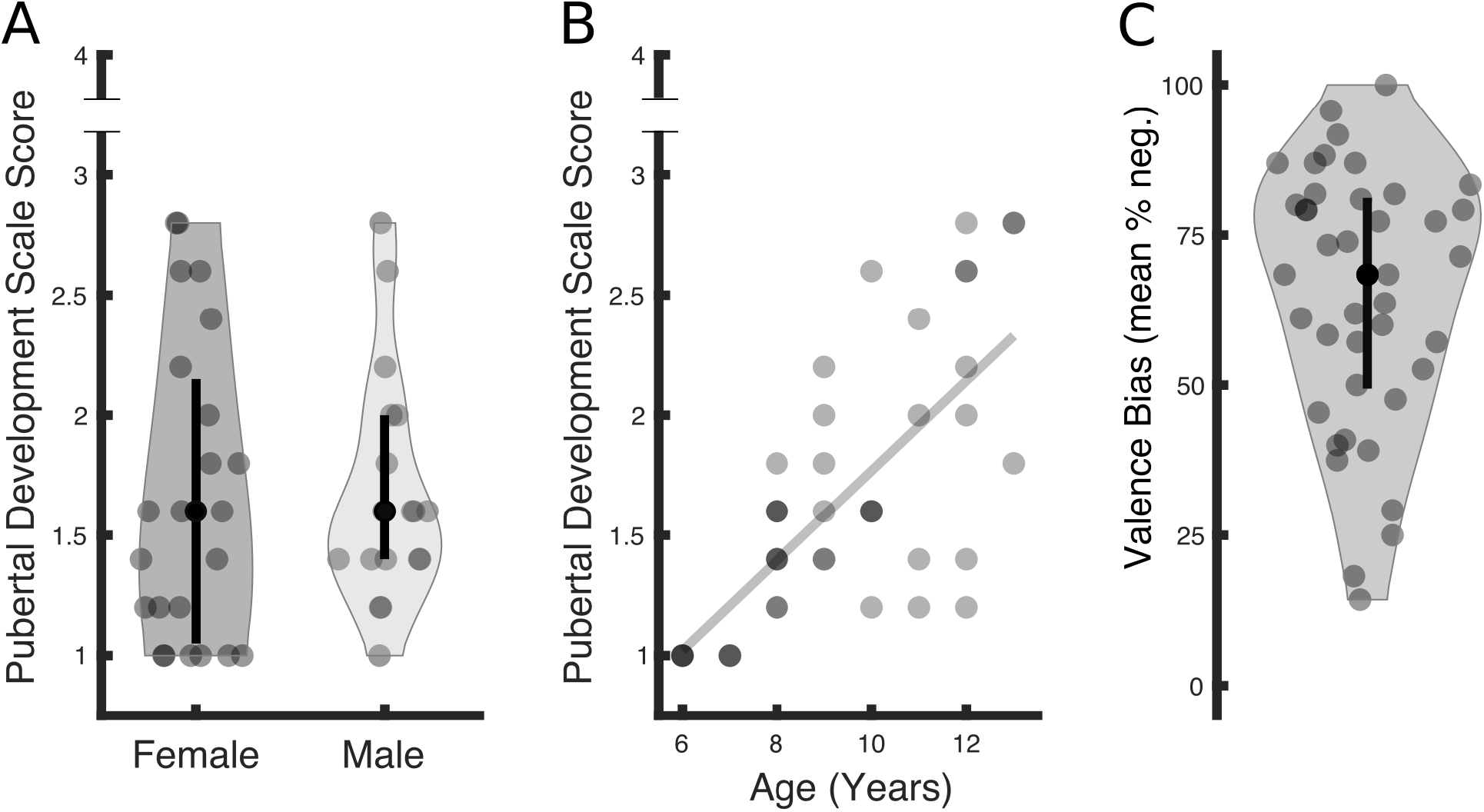
Descriptive illustration of measures. **(A)** The sample’s PDS scores ranged from 1 to 2.8, out of a highest possible score of 4. These scores subtend to a low range (mean(SD) = 1.67(.56), skewness(SD) = 0.71(0.37), kurtosis(SD) = −.54(.72)), consistent with the restricted age range (6-13 years) of the sample. Females (N = 23) and males (N = 18) did not differ as a function of PDS score (*t_39_* = 0.16, *p* = .87). In each violin plot, the black circle indicates the median (females = 1.60, males = 1.60), and the lower and upper ends of the black line indicate the first (females = 1.05, males = 1.40) and third (females = 2.15, males = 2.00) quartile cut-off, respectively. **(B)** Age was positively related to PDS score (*t*_39_ = 6.68, *p* < .001). Darker dots represent an overlap of participants with identical age/PDS values. **(C)** Valence bias scores, computed as the percent of negative ratings of surprised expressions, ranged from 14.29% to 100% with a mean of 64.66%. The black circle indicates the median (68.42%), and the lower and upper ends of the black line indicate the first (49.41%) and third (81.12%) quartile cut-offs, respectively.

Depressive symptoms were quantified using the major depression subscale of the Revised Child Anxiety and Depression Scale (RCADS; range = 0-12, possible range = 0-30, mean(SD) = 2.20(2.40); Chorpita, Yim, Moffitt, Umemoto, & Francis, 2000) administered to the child’s parent to maintain consistency across the sample given the young ages. This scale has been validated in ages as young as 3 years (Ebesutani et al., 2015). However, because our age range (6-13 years) extends below the minimum of standardized T scores (grades 3-12), raw scores of the major depression subscale were used in the analyses, consistent with a recent study (Vantieghem et al., 2017). One participant reported a major depression score of 12 which, while the age of this participant (7 years) was below the range included in the published T scores, was associated with a T score in the youngest standardized sample (3^rd^ grade) of 74, exceeding the cut-off associated with clinical diagnoses (T = 70; Chorpita et al., 2000). However, the inclusion of this participant did not qualitatively affect the reported results (see sections 3.1.2 and 3.2.1).

### 2.2 Procedure

#### 2.2.1 Session 1: behavioral

All experimental stimuli were presented on E-Prime software (version 2.0.10; Psychology Software Tools, Pittsburgh, PA). In Session 1 (Figure 2), participants performed a task to assess their baseline valence bias in which they viewed positive (e.g., happy), negative (e.g., angry), and ambiguous (e.g., surprised) images on a black background. For each image, participants were asked to make a two-alternative, forced-choice decision (via keyboard press) as quickly and as accurately as possible, indicating whether each image “felt good or felt bad”; this language and stimuli have been in previous work within similar age groups (Kestenbaum, 1992; Tottenham et al., 2013). This included a set of 48 faces, 24 with an ambiguous valence (surprised expression), and 24 with a clear valence (12 angry and 12 happy expressions). Of the 48 faces, 34 discrete face identities were used (17 males, 17 females) posing angry, happy, and surprised expressions. Fourteen identities (seven females, ages 21-30 years) were taken from the NimStim Set of Facial Expressions (Tottenham et al., 2009) and 20 (10 female, age 20-30 years) from the Karolinska Directed Emotional Faces database (Goeleven et al., 2008). All identities were European/European American. In separate but alternating blocks, scenes from the International Affective Picture System (IAPS; Lang, Bradley, & Cuthbert, 1997) were presented, consisting of 24 scenes with an ambiguous valence and 24 with a clear valence (12 negative and 12 positive). The purpose of rating the IAPS scenes was outside the scope of the current report, and did not contribute to the valence bias score nor were they included in any analysis. All blocks of stimuli included 24 images (12 ambiguous, 6 positive, and 6 negative) presented in a pseudorandom order (see Figure 2 for a depiction of tasks). Each stimulus was presented for 1500ms followed by a fixation cross for either 200 or 1900ms. We calculated the valence bias for each participant as the percentage of times that a participant indicated that a particular surprised face felt bad (e.g., a valence bias was 100% if that participant rated surprised faces as bad on all trials). Note that only the ratings of the surprised faces were used to calculate each valence bias score. Participants and parents then completed surveys (see above) and participants completed a mock scan (see below).

**Figure 2.**
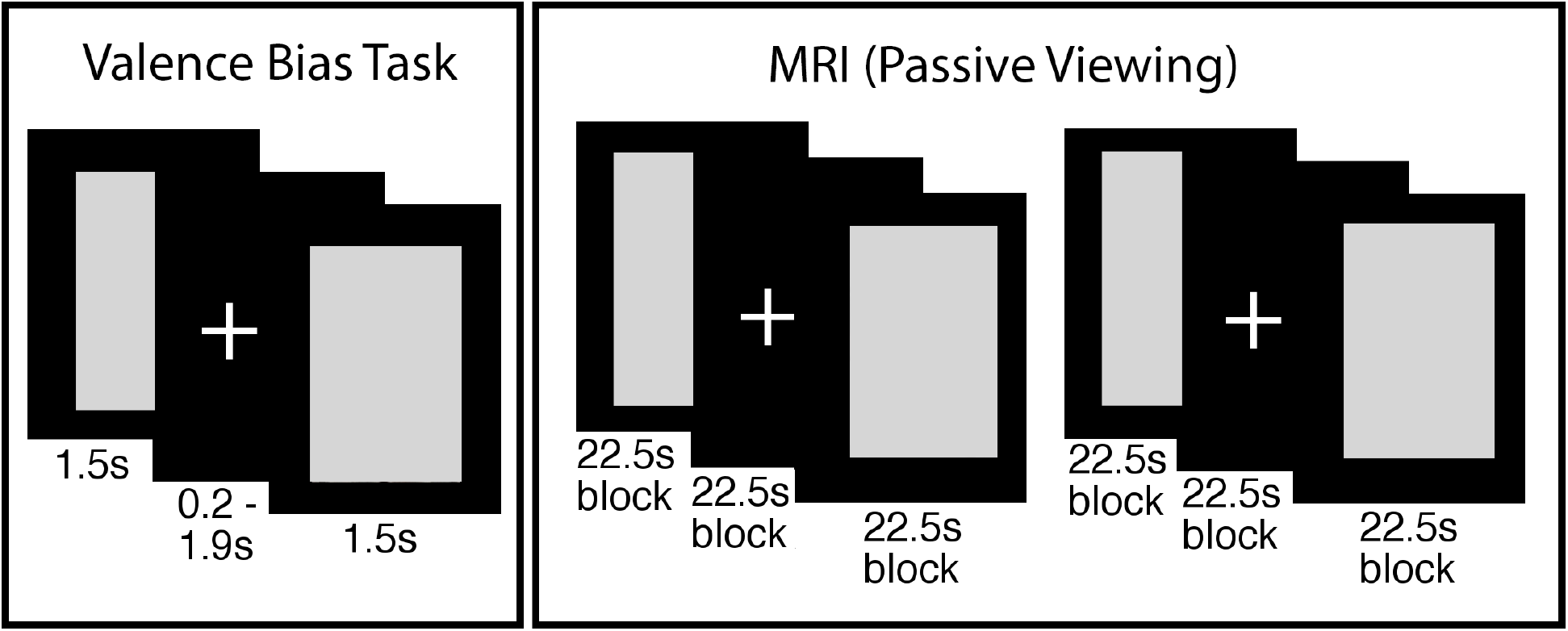
Depiction of procedure. In this illustration, gray boxes replace face stimuli as per bioRxiv requirements. In the valence bias task, participants viewed happy, angry, and surprised faces, and indicated whether each image “felt good or felt bad.” In the MRI, participants passively viewed new faces (i.e., not overlapping with the valence bias task) during two runs with blocks of surprised and neutral faces. Two runs with blocks of fearful and neutral faces followed, but were not included in the present analysis. Faces are from the NimStim Set of Facial Expressions (Tottenham et al., 2009).

#### 2.2.2 Session 2: MRI

Session 2 followed Session 1 by approximately 9 days (mean(SD) days = 9.34(6.69), range 1-46 days; Table 1). Days between sessions was unrelated to the behavioral (see section 3.1.2) and neuroimaging results (see section 3.2.1). Participants viewed a new set of European/European American faces from the Umeå University Database of Facial Expressions (Samuelsson et al., 2012) across 4 experimental runs while positioned in a MRI scanner (Figure 2). Prior to entering the MRI, all participants underwent a mock scanning session to acclimate to the environment and practice instructions to remain still during scanning. Padding was used to secure the participants’ head in a comfortable, static position. The experimenters provided feedback and reminders to remain still throughout the session.

**Table 1.** Descriptive statistics for behavioral variables.

| Variables | Mean (SD) | Min | Max | Min<br>Possible | Max<br>Possible |
| --- | --- | --- | --- | --- | --- |
| Valence Bias (mean % negative) | 64.66% (21.76%) | 14.29% | 100% | 0% | 100% |
| Depressive Symptoms (RCADS – Major Depression (Chorpita et al., 2000)) | 2.20 (2.40) | 0 | 12 | 0 | 30 |
| Petersen Pubertal Development Scale (PDS) score (Petersen et al., 1988) | 1.67 (.56) | 1 | 2.8 | 1 | 4 |
| Days Between Sessions | 9.34 (6.69) | 1 | 46 | — | — |

Each experimental run consisted of 4 blocks of 15 image presentations. Faces were presented for 500 ms and separated by a fixed interstimulus interval of 1000 ms. The first two runs consisted of 2 blocks of surprised faces and 2 blocks of neutral faces. Then, two additional runs followed which contained fearful instead of surprised faces, but their analysis was outside the scope of the present report. For the purpose of monitoring participants’ attention to the stimuli, participants were instructed to make a button response each time a flower appeared (instead of a face) on the screen; within each block, 3 images of flowers were pseudo-randomly presented amid the presentation of 12 face stimuli. All stimuli during this session were 500 x 750 pixels with a black background. Note that, consistent with extant work measuring valence bias, we used angry and happy faces in the behavioral task because they are clearly negative and positive and also perceptually distinct from surprised faces. However, for the MRI session, we used fearful and surprised faces because both recruit a similar dorsal region of the amygdala due to their increased uncertainty (Whalen, 2007) and because fearful faces are commonly used in studies measuring amygdala reactivity in development (Gee, Gabard-Durnam, et al., 2013; Gee, Humphreys, et al., 2013; Perlman & Pelphrey, 2011). However the scope of the current report is focused on the neural responses to surprised faces given their dual-valence ambiguity and use in the valence bias task.

### 2.3 MRI Parameters

#### 2.3.1 MRI data acquisition parameters

The MRI data were collected at the University of Nebraska-Lincoln, Center for Brain, Biology, & Behavior, on a Siemens 3T Skyra scanner using a 32-channel head coil. Structural images were acquired using a T1-weighted MPRAGE sequence (TR = 2.20 s, TE = 3.37 ms, slices = 192 interleaved, voxel size = 1.0 × 1.0 × 1.0 mm matrix = 256 × 256, FOV = 256 mm, flip angle = 7°, total acquisition time = 5:07). Blood oxygen level-dependent (BOLD) activity was measured using an EPI scanning sequence (TR = 2.50 s, TE = 30 ms, slides = 42 interleaved, voxel size = 2.50 × 2.50 × 2.80 mm, matrix = 88 × 88 mm, FOV = 220 mm, flip angle = 80°, total acquisition time = 3:14 per block) in which slices were acquired parallel to the intercommissural plane and the volume positioned to cover the extent of the entire brain.

#### 2.3.2 Preprocessing

Preprocessing of MR images was conducted using the Analysis of Functional Neuroimages (AFNI) program suite (Cox, 1996). The first four TRs of each run were excluded to allow for scanner stabilization. Voxel time-series were first despiked by removing values with outlying data in each separate voxel’s time-series. Then, slice timing correction was accomplished by re-referencing each scan to the first slice. These slice time corrected volumes were realigned to the minimum outlier image. Each volume was then aligned to the anatomical image before being warped to the Talairach template atlas (Talairach & Tournoux, 1988) provided by AFNI. Functional volumes were then spatially smoothed using a 6mm^3^ full width at half maximum kernel. The BOLD time-series, in each voxel separately, was normalized by dividing each time point by its average across all time points and then multiplying each time point by 100. Any images containing movements > .9mm^3^, as determined by the motion parameters calculated during spatial realignment, were censored frame-wise from further analysis. No participants showed excessive movements in more than 16% of their scans (mean(SD) = 2.47%(4.37%)), consistent with previous work (Gee, Humphreys, et al., 2013). The number of censored scans did not differ between surprise and neutral trials (*t*_40_ = 0.74, *p* = .46), and was not related to participant age (*t*_39_ = −1.50, *p* = .14).

### 2.4 Statistical Analysis

#### 2.4.1 Behavioral

See Table 1 for descriptive information of analyzed behavioral variables. Although the current study aimed to explore a puberty-moderated link between valence bias and both depressive symptoms and amygdala-mPFC connectivity, the crosssectional design limited the possibility of inferring directional associations between these variables (e.g., valence bias impacts depressive symptoms or depressive symptoms impacts valence bias). We conducted two separate moderation analyses, first within the behavioral data, and second related to the MRI data (see section 2.4.4).

First, to determine whether a more negative valence bias was related to higher depressive symptoms, the two measures were submitted to a robust bivariate regression (calculated using the *fitlm* command in Matlab) in which valence bias was a predictor of depression. Then, the moderating effect of pubertal status on the relationship between valence bias and depressive symptoms was tested in a robust regression with valence bias as the outcome variable; this model had a constant term and four predictors: 1) depressive symptoms, 2) PDS score, 3) the interaction between depressive symptoms × PDS score, and 4) age, which was included as a covariate. The interaction term coefficient represented the moderation effect. Importantly, because age was included as a covariate, the pubertal moderation results reflect the unique influence of this pubertal development score (above and beyond effects of age) on the relationship between valence bias and depressive symptoms.

Previous studies have related the age of pubertal onset with differences in emotion regulation and mental health outcomes (Gee, Gabard-Durnam, et al., 2013; Vantieghem et al., 2017). One possibility is that a moderation effect including valence bias and puberty could reflect a developmental “mismatch” between puberty and age, such that those who have a more negative valence bias are those who are either more or less developmentally mature than is typical for that age. To explore this possibility we calculated the “mismatch” between PDS score and age as the residuals of their relationship. We then submitted these residual scores (mean(SD) = 0.00(1.51), range = −1.94-3.78) to a robust regression with valence bias.

#### 2.4.2 BOLD reactivity

To identify brain regions in which BOLD activity related to each stimulus condition, the BOLD data were submitted to a general linear model (GLM) in which surprise, fear, and neutral blocks were modeled separately for the 4 runs of the experiment. These regressors included a boxcar function modeling the onset and duration of each block that was convolved with the hemodynamic response function. Each run included a constant term and nuisance regressors: 6 motion parameters (three rotational and three translational vectors) calculated during realignment, and both a linear and cubic polynomial trend model to control for BOLD signal drifts, consistent with AFNI guidelines given each runs’ 172.5 s duration.

#### 2.4.3 BOLD context-dependent connectivity

To assess the functional connectivity between the amygdala and the rest of the brain specific to each condition, a context-dependent connectivity analysis (i.e., psychophysiological interaction or PPI) was conducted using AFNI commands. The seed amygdala region was defined as the voxels showing activation while viewing surprised expressions compared to baseline (the fixation period between blocks). This contrast was used to identify all voxels sensitive to the surprised expressions. The resulting amygdala clusters were considered significant if exceeding a corrected threshold (FWE: *p* < .05) based on Gaussian Random Field theory (Friston et al., 1994; Hayasaka & Nichols, 2003) that avoids the spatial autocorrelation issue raised by Eklund and colleagues (Eklund et al., 2016). This threshold consisted of both a cluster-forming (*p* < .001) and cluster-extent (*k* > 21) threshold. One voxel cluster showed peak activation in the right basal forebrain (peak-*t*_40_ = 7.01, *k* = 86; x = 16, y = −4, z = −11) and extended into the dorsal amygdala (Figure 3A). The dorsal position of this cluster within the amygdala is consistent with previous work demonstrating that content conveying ambiguous valence recruits the amygdala/substantia innominata in particular (Kim, Somerville, McLean, et al., 2003; Whalen et al., 2009) rather than the more ventral regions that are relatively more sensitive to clear valence, such as expressions of anger (Whalen et al., 2001).

**Figure 3.**
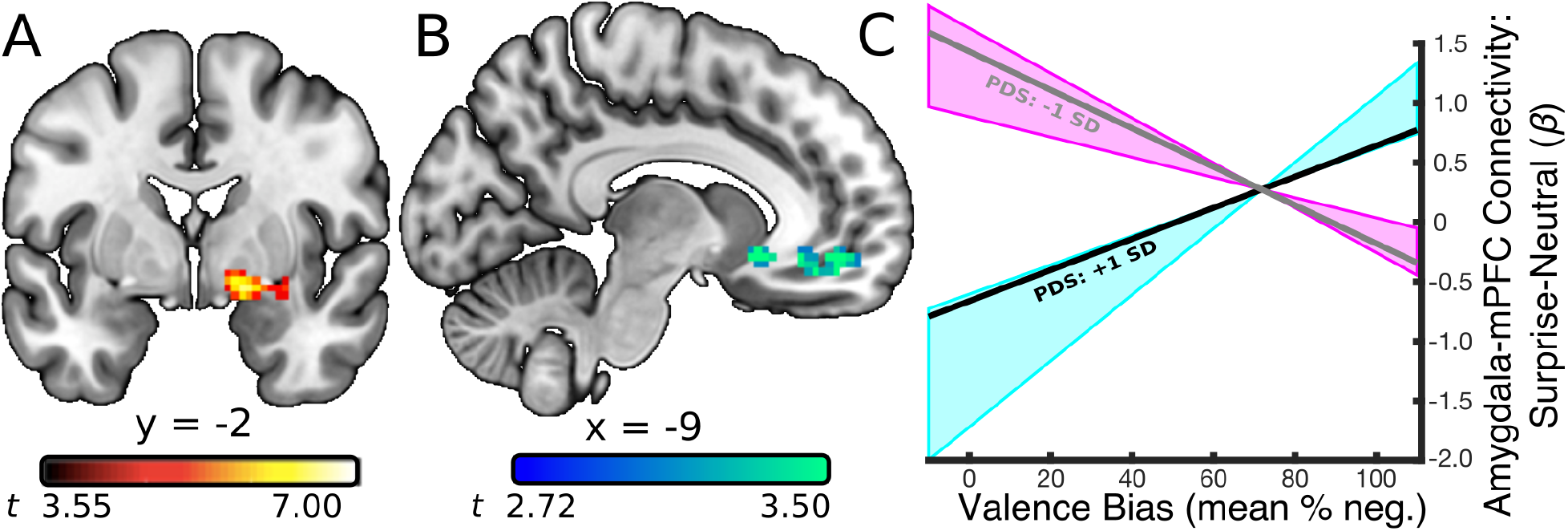
Relationship between amygdala-mPFC connectivity and negative valence bias as a function of PDS score. **A)** A seed region in the right amygdala was defined using the contrast of surprised facial expressions versus baseline (*p* < 0.0005). The dorsal position of this cluster within the amygdala is consistent with previous work demonstrating that content conveying ambiguous valence recruits the amygdala/substantia innominata in particular (Kim, Somerville, McLean, et al., 2003; Whalen et al., 2009). **B)** A PPI analysis based on surprise > neutral activity in the amygdala seed revealed a relationship between amygdala connectivity and valence bias that was moderated by PDS score in the mPFC (peak-*t*_36_ = 4.50, *p* = .00007). **C)** The estimated regression slopes between valence bias and the surprise > neutral amygdala-mPFC connectivity betas, at a lower (gray line; 1 standard deviation below the mean PDS score; *t*_36_ = −4.89, *p* = .00002) and relatively higher (black line; 1 standard deviation above the mean PDS score; *t*_36_ = 3.52, *p* = .001) PDS scores. More mature children (within this relatively immature sample; blue shaded area denotes moderator’s region of significance) that have a mature (inverse) connectivity pattern were more likely to have a positive valence bias while those that have the less mature (positive) connectivity pattern were more likely to show a negative valence bias. Lower PDS scores (pink shaded area denotes moderator’s region of significance) predicted the *opposite* relationship between valence bias and amygdala-mPFC connectivity.

To model the face-evoked BOLD activity in this amygdala region, the BOLD activity from this dorsal amygdala region was first deconvolved with a hemodynamic response function, and then multiplied with boxcar functions modeling stimulus onsets and durations separately for each condition, resulting in 4 condition-specific models of amygdala activity. Lastly, these condition-specific amygdala regressors were convolved with a hemodynamic response function. These regressors entered a GLM with a constant term, task onset regressors, and a model of the amygdala activity across the whole duration of the experiment. Thus, the beta values associated with the condition-specific amygdala regressors reflect changes in amygdala connectivity evoked by the blocked stimulus presentation. As in the previously described analysis of BOLD reactivity, nuisance regressors consisted of the 6 motion models to control for movement artifacts and, because the BOLD activity was analyzed across all runs continuously (690 seconds), 5 polynomial trends to control for BOLD signal drifts, consistent with AFNI guidelines.

#### 2.4.4 mPFC-amygdala BOLD connectivity and valence bias

Connectivity between the amygdala and mPFC during emotional processing changes across development, such that there is a shift toward more inverse connectivity around age 10 years, thought to support age-related changes in emotion regulatory behaviors (Gee, Humphreys, et al., 2013). The amygdala seed region used in the current connectivity analysis was defined as voxels showing significant activation for surprise relative to baseline, a criterion chosen in order to include as many voxels sensitive to the surprised face expressions as possible. For comparison with valence bias and pubertal status, the connectivity beta differences for surprise *relative to neutral* were used in order to isolate amygdala connectivity specific to the ambiguity conveyed through the surprised expressions rather than a general response to facial expressions. Thus, these surprise > neutral connectivity beta differences for each participant were submitted to a robust multiple regression (calculated using the *fitlm* command in Matlab), separately at each voxel, with a constant term and four predictors: 1) valence bias, 2) PDS score, 3) the interaction between valence bias × PDS score, and 4) age, which was included as a covariate. To determine whether pubertal status moderated the relationship between valence bias and amygdala connectivity in any mPFC region, the interaction term coefficients, computed at each voxel separately, were passed through a cluster-forming (*p* < .01) and -extent threshold (*k* > 75) according to Guassian Random Field theory guidelines for multiple comparison correction (Friston et al., 1994; Hayasaka & Nichols, 2003). Importantly, as in the moderation analysis of behavior (see section 2.4.1), because age was included as a covariate, the pubertal moderation results reflect the unique influence of puberty (above and beyond the effects of age) on the relationship between valence bias and amygdala connectivity.

#### 2.4.5 Individual differences between BOLD, age, and puberty

Lastly, we examined the direct relationship between the BOLD measures (amygdala activation and connectivity with mPFC) and age and pubertal status. Specifically, we calculated the average betas values, separately for each participant, for 1) the surprise > neutral BOLD activation for all voxels in the amygdala seed (see section 2.4.3) and 2) the surprise > neutral amygdala connectivity for all voxels in the mPFC (see section 3.2.1). These average beta values were submitted to a robust regression with both age and PDS score, yielding four separate beta coefficients.

## 3. Results

### 3.1 Behavioral

#### 3.1.1 Valence ratings – characterizing valence bias

Participants rated angry faces as negative (mean(SD) % negative = 85.91(11.75); range = 62-100), and happy faces as positive (mean(SD) % negative = 10.18(9.10); range = 0-30). In contrast, ratings of surprised faces showed greater variability (Figure 1C; mean(SD) % negative = 64.66(21.75); range = 14.29-100), and represented the valence bias for each individual, such that higher scores indicated a more negative bias. Within this restricted age range, valence bias was not significantly related to age (*t*_39_ = −0.48, *p* = .63) or PDS score (*t*_39_ = −0.42, *p* = .68).

#### 3.1.2 Depressive symptoms and valence bias moderated by pubertal status

Table 1 lists the descriptive statistics for the behavioral variables. A bivariate robust regression revealed that valence bias was positively related to depressive symptoms (*t*_39_ = 2.29, *p* = .03), such that those with a more negative bias had higher depressive symptoms. For the moderation analysis, the interaction between depressive symptoms and PDS score was significant (*t*_36_ = 2.48, *p* = .02; Figure 4) indicating that pubertal status, when controlling for age, moderated the relationship between depression and valence bias.

**Figure 4.**
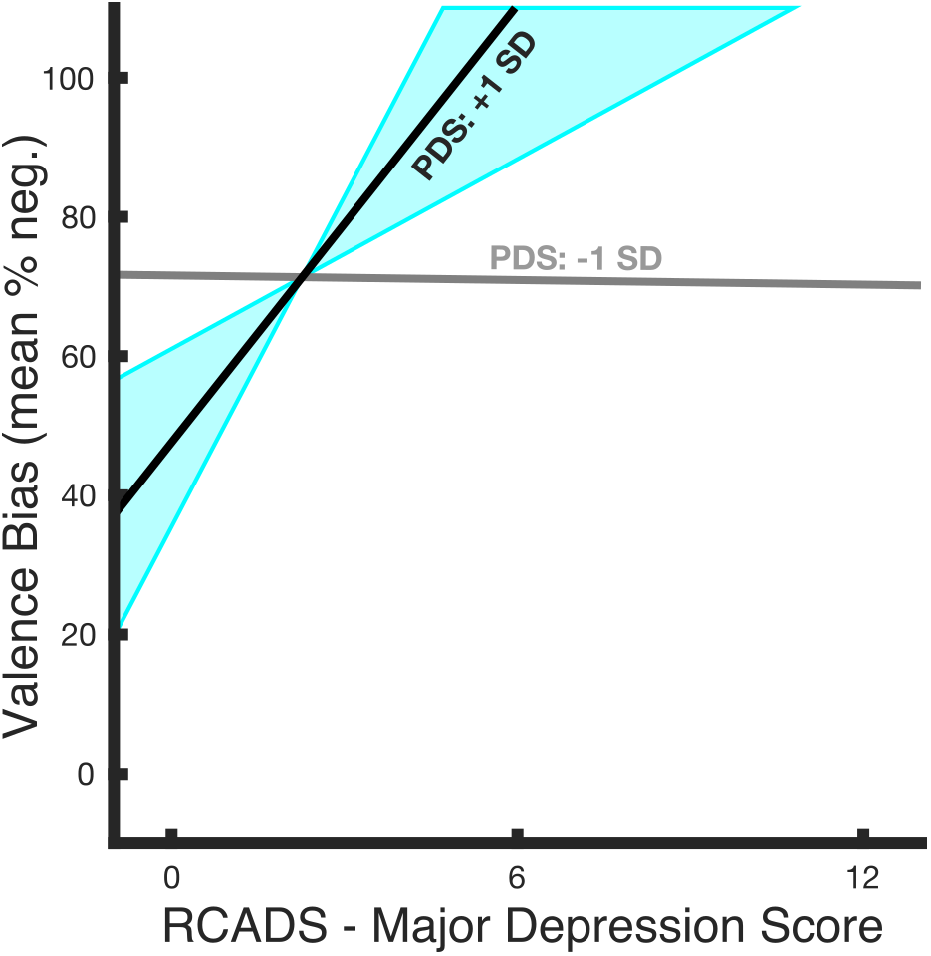
Relationship between valence bias and depressive symptoms as a function of PDS score. The relationship between depressive symptoms and valence bias was moderated by PDS score (*t_36_* = 2.48, *p* = .02) such that valence bias and depressive symptoms shared a stronger positive relationship at relatively higher PDS scores. The estimated regression slopes for the relationship between valence bias and depressive symptoms at different PDS scores illustrate that a lower PDS score (gray line; 1 standard deviation below the mean PDS score; *t*_36_ = −.06, *p* = .95) predicted no relationship, whereas a higher PDS score (black line; 1 standard deviation above the mean pubertal status; *t*_36_ = 2.63, *p* = .01) predicted a positive relationship (blue shaded area denotes moderator’s region of significance). At no point did lower PDS scores predict a significant relationship between valence bias and depressive symptoms, therefore no shaded area (region of significance) is illustrated around the −1 SD line.

To explore this moderation, the relationship between valence bias and depressive symptoms was calculated for each PDS score. This relationship became significant between a PDS score of 1.6-2.8 (out of the sample’s range of 1-2.8; Figure 4, blue shaded area). To test for potential confounding effects, we ran similar moderation analyses that included sex (male or female), or number of days between sessions 1 and 2, or excluded the participant with a suprathreshold clinical major depression score, or excluding participants who did not identify as Caucasian. Notably, the effects were qualitatively the same with each of these modifications, suggesting these variables did not impact the reported findings.

As an additional analysis we tested whether a developmental “mismatch” between puberty and age (e.g., an accelerated pubertal development) is associated with valence bias, which may potentially confound the moderation effects reported above. We calculated this “mismatch” between PDS score and age as the residuals of their relationship and found that this was not related to valence bias (*t*_39_ = −0.17, *p* = .86).

### 3.2 MRI

#### 3.2.1 Context-dependent amygdala-mPFC connectivity and valence bias

The relationship between surprise > neutral amygdala connectivity and valence bias, when controlling for age, was moderated by pubertal status in two clusters (Table 2). In other words, those with a higher PDS score (within this relatively immature sample) showed an effect whereby a more positive valence bias was associated with a more inverse (mature, i.e., suggestive of emotion regulation) connectivity in these regions. The first cluster showed peak activation in the right subcollosal gyrus (peak-*t*_36_ = 5.52, *p* = .000003; *k*=170; Talaraich (x, y, z) coordinates: 11, 11, −14) and extended into the right rectal gyrus and right anterior cingulate. The second cluster showed peak activation in the left medial frontal gyrus (Figure 3B; peak-*t*_36_ = 4.50, *p* = .00007; *k*=92; Talaraich (x, y, z) coordinates: −11, 49, −11) and extended into the left anterior cingulate, consistent with reports of amygdala connectivity with the mPFC (Kim, Somerville, Johnstone, et al., 2003).

**Table 2.**
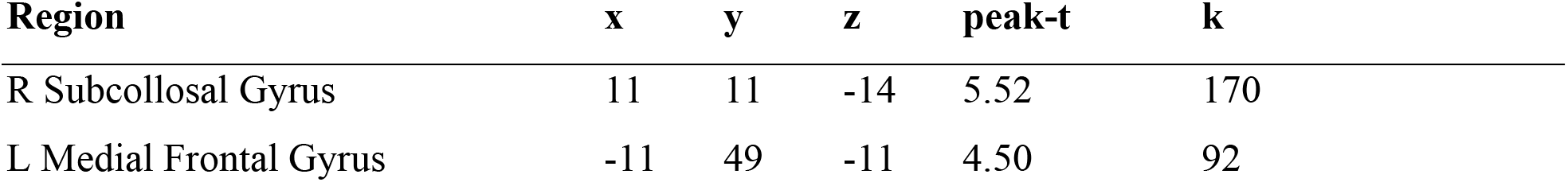
Clusters of significant BOLD surprise > neutral amygdala connectivity whose relationship with valence bias was moderated by pubertal score

To explore the moderation within this latter mPFC cluster, the conditional effects of pubertal status were calculated from the intercept and slope of the relationship between valence bias and the voxel averaged amygdala-mPFC connectivity beta values; this was done separately at each PDS score in this sample (range = 1-2.8). These conditional effects (Figure 3C) indicated that for those with a relatively higher PDS score (2-2.8; i.e., the highest scores in this sample), the relationship between valence bias and amygdala-mPFC connectivity was more positive (Figure 3C, blue shaded area). In other words, children with both a higher PDS score and a less mature (positive) connectivity pattern had a more negative valence bias. In contrast, children with both a higher PDS score and a more mature (inverse) connectivity pattern had a more positive valence bias. The opposite effect (i.e., a more negative valence bias predicted more inverse amygdala-mPFC connectivity) was significant at PDS scores 1-1.4 (Figure 3C, pink shaded area).

Finally, as with the behavioral results (see section 3.1.2), we tested for potential confounding effects by rerunning the moderation analyses and including sex, or number of days between sessions 1 and 2, or excluding the participant with a supra-threshold clinical major depression score, or excluding participants who did not identify as Caucasian. In addition, although surprise > neutral amygdala activation was not related to amygdala connectivity (*t*_39_ = - 1.10, *p* = .28), we ran another moderation which included amygdala activation as a covariate to test whether the connectivity moderation effect was confounded by the level of BOLD activation. Notably, the effects were qualitatively the same with each of these modifications, suggesting these variables did not impact the reported findings.

#### 3.2.2 Individual differences between BOLD, age, and puberty

To describe the basic relationship between the BOLD measures and age and puberty, the surprise > neutral amygdala BOLD activation betas and the amygdala-mPFC connectivity betas were submitted to a robust regression with age and PDS score. Neither amygdala activation nor connectivity were related to age (activation: *t*_39_ = 0.68, *p* = .50, connectivity: *t*_39_ = −0.19, *p* = .85) or PDS score (activation: *t*_39_ = 0.08, *p* = .94, connectivity: *t*_39_ = −0.37, *p* = .72).

### 4. Discussion

Within this relatively pubertally immature sample, higher PDS scores coupled with a more negative valence bias were associated with more depressive symptoms and less inverse amygdala-mPFC connectivity (suggestive of weaker emotion regulation). Broadly, these exploratory results support the notion that internalizing problems, such as depression, arise from dysfunctional emotion regulation circuitry (Banks et al., 2007; Phillips et al., 2008) and, more specifically, that this link arises from developmental differences (Gee, Gabard-Durnam, et al., 2013; Gee, Humphreys, et al., 2013; Hare et al., 2008; Perlman & Pelphrey, 2011). Given that this amygdala-mPFC circuit is intimately tied to biological changes during puberty (Andersen & Teicher, 2008; Angold & Costello, 2006; Paus et al., 2008), the early pubertal period explored in the current study may be critical in the construction of healthy emotion regulation mechanisms. These exploratory results support a model for future research which predicts that, while negative valence bias in early childhood is normative, negative bias in later development is putatively *maintained* by the failure to develop a more mature, regulatory frontoamygdalar circuitry, which may *increase the risk* for depression.

These findings are consistent with our “initial negativity hypothesis” that posits that the initial or default interpretation of surprise is more negative (Neta et al., 2011; Neta & Tong, 2016; Neta & Whalen, 2010). In contrast, positive ratings depend upon slower and more elaborate emotion regulation processes which override the initial negativity and putatively downregulate the amygdala response (Kaffenberger et al., 2010; Kim, Somerville, Johnstone, et al., 2003; Neta et al., 2011; Neta & Tong, 2016; Neta & Whalen, 2010; Petro et al., 2018), processes that are likely compromised in depression and anxiety (Beck, 1976; Reef et al., 2011; Williams et al., 2007). Indeed, age-related differences in this emotion regulation circuity (i.e., amygdala-mPFC connectivity (Gee, Humphreys, et al., 2013)) are associated with mental health risk factors in adults (Hare et al., 2008). The current results extend this “initial negativity hypothesis” by suggesting that individual differences in valence bias may originate during pubertal development, alongside the development of this emotion regulatory circuitry that putatively overrides the default, or initial negativity.

The utility of surprised faces in tracking individual differences in negativity bias and emotion regulation brain circuits is broadly consistent with a functional – contextual account of facial displays (Crivelli & Fridlund, 2018), which predicts that an expression’s emotional value depends on its social, environmental, and/or cultural context (Barrett & Kensinger, 2010). Whereas happy and angry expressions signal predominantly positive or negative social outcomes, respectively, surprised faces signal multiple possible outcomes spanning positive and negative valence (i.e., dual-valence ambiguity). Thus, the valence assigned to surprised expressions presented without context is more pliable to interpretation. The use of stimuli with dual-valence ambiguity represents a methodological advance in conceptualizing negativity bias in that it side-steps limitations present in extant literature. For instance, negativity bias is often measured via either an attentional bias toward clearly negative *or* away from clearly positive stimuli (see Fales et al., 2008) or by examining responses to ambiguous stimuli with alternate meanings that could be either negative or neutral, but not positive (e.g. “lie”; Mathews et al., 1989). These findings not only rely on cognitive/linguistic abilities not developmentally appropriate for children, but by not examining responses to stimuli with a dual-valence representation (i.e., negative *and* positive possible interpretations), these earlier findings are skewed toward the extremes of the valence spectrum. As such, this earlier work is limited in its ability to identify individual differences in responses to emotional stimuli during sensitive periods of development, and thus has more limited findings regarding the developmental origins of negativity bias. Future work will benefit from incorporating our valence bias task to examine individual differences in emotion reactivity and regulation, particularly in young ages.

These findings make a meaningful first step toward establishing the developmental origins of negativity bias. One caveat is the relatively small sample size with somewhat limited variance in depressive symptoms, and that the PDS was only administered to children ages 8 years and older. Future work should replicate our exploratory findings in larger samples with a wider range of depressive symptoms and also determine the extent to which these findings, which use parent-reported subclinical depression (see; Muris, Meesters, & Fijen, 2003), generalize to those with clinically diagnosed or self-reported depression. Further, future work should also explore the extent to which the effects generalize to the range of mental health disorders in which internalizing problems manifest. Finally, although there was no explicit emotion regulation task, a more inverse frontoamygdalar connectivity is thought to represent emotion regulation (Gee, Humphreys, et al., 2013; Ochsner et al., 2004). Indeed, the connectivity in these children spanned both positive and negative values, consistent with previous studies (Gee, Humphreys, et al., 2013; Hare et al., 2008) which suggest that *inverse* connectivity (rather than a *lack of positivity connectivity*) may be associated with a more adult-like regulatory circuitry. While the current study treated connectivity as a continuous measure in order to test its relationship to valence bias and pubertal status, future work with larger sample sizes in normative adults should aim to identify the point at which negative frontoamygdalar connectivity defines an emotion regulatory process.

Whether or not long-term mental health trajectories are impacted by environmental factors may also be explored in future research. For instance, early life stress is a risk factor for mental health disorders (Tottenham et al., 2011) and is associated with developmental differences in regulatory circuitry (Cohodes et al., 2020; Heim & Binder, 2012; Lupien et al., 2009). Recent work suggests that positive affect may serve as a buffer from the development of depressive symptoms (Sewart et al., 2019), such that the risk for internalizing problems imposed by early life stress is mitigated by an accelerated onset of puberty which leads to an earlier developmental shift towards positive valence bias (Gee, Gabard-Durnam, et al., 2013; Vantieghem et al., 2017). This suggests that resiliency, or the ability to find a positive outlook in potentially negative situations (Gross & John, 2003), implicates the same emotion regulation mechanism explored in the current study (Tugade & Fredrickson, 2004). In contrast to this prediction that an accelerated onset of puberty is related to an earlier development of emotion regulation skills, those with higher PDS scores given their age in the current study’s sample were not more positive. However, future studies may further explore whether or not normative puberty-by-age differences predict differences in valence bias and emotion regulation skills.

The moderating role of puberty on the relationship between valence bias and depressive symptoms occurred earlier in maturation (starting at a PDS score of 1.6 and up to the highest score in this sample of 2.8) than the relationship between valence bias and brain connectivity (starting at a PDS score of 2 and up to the highest score in this sample of 2.8). Although crosssectional, these findings provide preliminary evidence that a change toward a more positive valence bias and away from depressive symptoms may have downstream effects on developing more adult-like brain connectivity patterns. Future longitudinal work should extend our exploratory work and aim to 1) test the prediction that the *maintenance* of negative valence bias is associated with both an increase in depressive symptoms and a slower development of (inverse) amygdala-mPFC connectivity, and 2) establish the directionality these relationships. The opposite relationship between valence bias and amygdala-mPFC connectivity was also observed at a PDS score of 1.4 and below. However, our predictions about the effects in those at the earliest stages of puberty were less clear, particularly before the point at which puberty moderates the link between valence bias and depressive symptoms (below a PDS score of 1.6) and given the assumed scores in children ages 6-7 years).

Because females and males show different age onsets in puberty (Schuiling & Likis, 2016), maturity was measured via a scale of pubertal development. In addition, follow-up analyses found sex was not related to the pubertal moderation of amygdala-mPFC connectivity. Considering the age range of the current sample (6-13 years), these results are consistent with the finding that increased depression in females emerges *after* the age of 13 (Ferguson et al., 1999; Hankin & Abramson, 2001). Continued work should explore whether sex differences emerge in older ages.

Future longitudinal work may also hold broad implications for treating mental health. For instance, the effects reported here may pinpoint developmental periods most sensitive to longterm mental health outcomes. Such information will be critical for informing potential interventions (e.g., mindfulness), which can improve emotion regulation success and decrease negativity bias (Goldin & Gross, 2010; Jazaieri et al., 2014). These types of training may be particularly useful for individuals that putatively maintain a negativity bias beyond a normative developmental stage such that this bias interferes with normal, healthy functioning.

## Acknowledgements

This work was supported by the National Institutes of Health (NIMH111640; PI: Neta), and by Nebraska Tobacco Settlement Biomedical Research Enhancement Funds. We thank Rebecca L. Brock for consultation regarding statistical analyses. We thank R. James R. Blair and Leah H. Somerville for helpful comments on the manuscript. We also thank Ruby Basyouni, Kayla Clark, Daniel J. Henley and Tien T. Tong for assistance in data collection and management.

## Competing Interests

The authors declare that they have no competing interests.

## Data Availability

The data relevant to this manuscript are available on the NIH National Database Archive.

## Notes

### Competing Interest Statement

The authors have declared no competing interest.

## References

Andersen, S. L., & Teicher, M. H. (2008). Stress, sensitive periods and maturational events in adolescent depression. Trends in Neurosciences, 31(4), 183–191. https://doi.org/10.1016/j.tins.2008.01.004

Angold, A., & Costello, E. J. (2006). Puberty and Depression. Child and Adolescent Psychiatric Clinics of North America, 15(4), 919–937. https://doi.org/10.1016/j.chc.2006.05.013

Asato, M. R., Terwilliger, R., Woo, J., & Luna, B. (2010). White Matter Development in Adolescence: A DTI Study. Cerebral Cortex, 20(9), 2122–2131. https://doi.org/10.1093/cercor/bhp282

Banks, S. J., Eddy, K. T., Angstadt, M., Nathan, P. J., & Phan, K. L. (2007). Amygdala–frontal connectivity during emotion regulation. Social Cognitive and Affective Neuroscience, 2(4), 303–312. https://doi.org/10.1093/scan/nsm029

Barrett, L. F., & Kensinger, E. A. (2010). Context is routinely encoded during emotion perception. Psychological Science, 21(4), 595–599. https://doi.org/10.1177/0956797610363547

Beck, A. T. (1976). Cognitive therapy and the emotional disorders. International Universities Press.

Blakemore, S.-J., Burnett, S., & Dahl, R. E. (2010). The role of puberty in the developing adolescent brain. Human Brain Mapping, 31(6), 926–933.

Blanton, R. E., Cooney, R. E., Joormann, J., Eugène, F., Glover, G. H., & Gotlib, I. H. (2012). Pubertal stage and brain anatomy in girls. Neuroscience, 217, 105–112. https://doi.org/10.1016/j.neuroscience.2012.04.059

Brown, C. C., Raio, C. M., & Neta, M. (2017). Cortisol responses enhance negative valence perception for ambiguous facial expressions. Scientific Reports, 7. https://doi.org/10.1038/s41598-017-14846-3

Browning, M., Holmes, E. A., & Harmer, C. J. (2010). The modification of attentional bias to emotional information: A review of the techniques, mechanisms, and relevance to emotional disorders. Cognitive, Affective, & Behavioral Neuroscience, 10(1), 8–20. https://doi.org/10.3758/CABN.10.1.8

Bruce, V., Campbell, R. N., Doherty-Sneddon, G., Langton, S., McAuley, S., & Wright, R. (2000). Testing face processing skills in children. British Journal of Developmental Psychology, 18(3), 319–333. https://doi.org/10.1348/026151000165715

Burghy, C. A., Stodola, D. E., Ruttle, P. L., Molloy, E. K., Armstrong, J. M., Oler, J. A., Fox, M. E., Hayes, A. S., Kalin, N. H., Essex, M. J., Davidson, R. J., & Birn, R. M. (2012). Developmental pathways to amygdala-prefrontal function and internalizing symptoms in adolescence. Nature Neuroscience, 15(12), 1736–1741. https://doi.org/10.1038/nn.3257

Burke, K. C. (1991). Comparing Age at Onset of Major Depression and Other Psychiatric Disorders by Birth Cohorts in Five US Community Populations. Archives of General Psychiatry, 48(9), 789. https://doi.org/10.1001/archpsyc.1991.01810330013002

Chorpita, B. F., Yim, L., Moffitt, C., Umemoto, L. A., & Francis, S. E. (2000). Assessment of symptoms of DSM-IV anxiety and depression in children: A revised child anxiety and depression scale. Behaviour Research and Therapy, 38(8), 835–855.

Clark, D. A., & Beck, A. T. (2010). Cognitive theory and therapy of anxiety and depression: Convergence with neurobiological findings. Trends in Cognitive Sciences, 14(9), 418–424. https://doi.org/10.1016/j.tics.2010.06.007

Cohodes, E. M., Kitt, E. R., Baskin-Sommers, A., & Gee, D. G. (2020). Influences of early-life stress on frontolimbic circuitry: Harnessing a dimensional approach to elucidate the effects of heterogeneity in stress exposure. Developmental Psychobiology. https://doi.org/10.1002/dev.21969

Cox, R. W. (1996). AFNI: Software for Analysis and Visualization of Functional Magnetic Resonance Neuroimages. Computers and Biomedical Research, 29(3), 162–173. https://doi.org/10.1006/cbmr.1996.0014

Crivelli, C., & Fridlund, A. J. (2018). Facial displays are tools for social influence. Trends in Cognitive Sciences, 22(5), 388–399. https://doi.org/10.1016/j.tics.2018.02.006

Das, P., Kemp, A. H., Flynn, G., Harris, A. W. F., Liddell, B. J., Whitford, T. J., Peduto, A., Gordon, E., & Williams, L. M. (2007). Functional disconnections in the direct and indirect amygdala pathways for fear processing in schizophrenia. Schizophrenia Research, 90(1), 284–294. https://doi.org/10.1016/j.schres.2006.11.023

Denham, S. A. (1998). Emotional development in young children. Guilford Press.

Dorn, L. D., Dahl, R. E., Woodward, H. R., & Biro, F. (2006). Defining the Boundaries of Early Adolescence: A User’s Guide to Assessing Pubertal Status and Pubertal Timing in Research With Adolescents. Applied Developmental Science, 10(1), 30–56. https://doi.org/10.1207/s1532480xads1001_3

Dubois, J., & Adolphs, R. (2016). Building a Science of Individual Differences from fMRI. Trends in Cognitive Sciences, 20(6), 425–443. https://doi.org/10.1016/j.tics.2016.03.014

Ebesutani, C., Tottenham, N., & Chorpita, B. (2015). The revised child anxiety and depression scale - parent version: Extended applicability and validity for use with younger youth and children with histories of early-life caregiver neglect. Journal of Psychopathology and Behavioral Assessment, 37(4), 705–718. https://doi.org/10.1007/s10862-015-9494-x

Eisenberg, N., Fabes, R. A., Bernzweig, J., Karbon, M., Poulin, R., & Hanish, L. (1993). The relations of emotionality and regulation to preschoolers’ social skills and sociometric status. Child Development, 64(5), 1418–1438.

Eklund, A., Nichols, T. E., & Knutsson, H. (2016). Cluster failure: Why fMRI inferences for spatial extent have inflated false-positive rates. Proceedings of the National Academy of Sciences of the United States of America, 113(28), 7900–7905. https://doi.org/10.1073/pnas.1602413113

Emslie, G. J., Mayes, T. L., & Ruberu, M. (2005). Continuation and Maintenance Therapy of Early-Onset Major Depressive Disorder: Pediatric Drugs, 7(4), 203–217. https://doi.org/10.2165/00148581-200507040-00001

Etkin, A., Prater, K. E., Schatzberg, A. F., Menon, V., & Greicius, M. D. (2009). Disrupted Amygdalar Subregion Functional Connectivity and Evidence of a Compensatory Network in Generalized Anxiety Disorder. Archives of General Psychiatry, 66(12), 1361. https://doi.org/10.1001/archgenpsychiatry.2009.104

Fales, C. L., Barch, D. M., Rundle, M. M., Mintun, M. A., Snyder, A. Z., Cohen, J. D., Mathews, J., & Sheline, Y. I. (2008). Altered emotional interference processing in affective and cognitive-control brain circuitry in major depression. Biological Psychiatry, 63(4), 377–384. https://doi.org/10.1016/j.biopsych.2007.06.012

Ferguson, T. J., Stegge, H., Miller, E. R., & Olsen, M. E. (1999). Guilt, shame, and symptoms in children. Developmental Psychology, 35(2), 347–357. https://doi.org/10.1037/0012-1649.35.2.347

Forbes, E. E., Phillips, M. L., Silk, J. S., Ryan, N. D., & Dahl, R. E. (2011). Neural Systems of Threat Processing in Adolescents: Role of Pubertal Maturation and Relation to Measures of Negative Affect. Developmental Neuropsychology, 36(4), 429–452. https://doi.org/10.1080/87565641.2010.550178

Friston, K. J., Worsley, K. J., Frackowiak, R. S., Mazziotta, J. C., & Evans, A. C. (1994). Assessing the significance of focal activations using their spatial extent. Human Brain Mapping, 1(3), 210–220. https://doi.org/10.1002/hbm.460010306

Gabard-Durnam, L. J., Flannery, J., Goff, B., Gee, D. G., Humphreys, K. L., Telzer, E., Hare, T., & Tottenham, N. (2014). The development of human amygdala functional connectivity at rest from 4 to 23 years: A cross-sectional study. NeuroImage, 95, 193–207. https://doi.org/10.1016/j.neuroimage.2014.03.038

Gee, D., Gabard-Durnam, L., Flannery, J., Goff, B., Humphreys, K., Telzer, E., Hare, T., Bookheimer, S., & Tottenham, N. (2013). Early developmental emergence of human amygdala–prefrontal connectivity after maternal deprivation. Proceedings of the National Academy of Sciences of the United States of America, 110(39), 15638–15643. https://doi.org/10.1073/pnas.1307893110

Gee, D., Humphreys, K., Flannery, J., Goff, B., Telzer, E. H., Shapiro, M., Hare, T., Bookheimer, S., & Tottenham, N. (2013). A developmental shift from positive to negative connectivity in human amygdala–prefrontal circuitry. Journal of Neuroscience, 33(10), 4584–4593. https://doi.org/10.1523/JNEUROSCI.3446-12.2013

Goddings, A., Heyes, S. B., Bird, G., Viner, R. M., & Blakemore, S. J. (2012). The relationship between puberty and social emotion processing. Developmental Science, 15(6), 801–811. https://doi.org/10.1111/j.1467-7687.2012.01174.x

Goddings, A.-L., Mills, K. L., Clasen, L. S., Giedd, J. N., Viner, R. M., & Blakemore, S.-J. (2014). The influence of puberty on subcortical brain development. NeuroImage, 88, 242–251. https://doi.org/10.1016/j.neuroimage.2013.09.073

Goeleven, E., De Raedt, R., Leyman, L., & Verschuere, B. (2008). The karolinska directed emotional faces: A validation study. Cognition and Emotion, 22(6), 1094–1118. https://doi.org/10.1080/02699930701626582

Goldin, P. R., & Gross, J. J. (2010). Effects of mindfulness-based stress reduction (mbsr) on emotion regulation in social anxiety disorder. Emotion, 10(1), 83–91. https://doi.org/10.1037/a0018441

Gross, J. J., & John, O. P. (2003). Individual differences in two emotion regulation processes: Implications for affect, relationships, and well-being. Journal of Personality and Social Psychology, 85(2), 348–362. https://doi.org/10.1037/0022-3514.85.2.348

Guyer, A. E., Monk, C. S., McClure-Tone, E. B., Nelson, E. E., Roberson-Nay, R., Adler, A. D., Fromm, S. J., Leibenluft, E., Pine, D. S., & Ernst, M. (2008). A developmental examination of amygdala response to facial expressions. Journal of Cognitive Neuroscience, 20(9), 1565–1582. https://doi.org/10.1162/jocn.2008.20114

Hankin, B. L., & Abramson, L. Y. (2001). Development of gender differences in depression: An elaborated cognitive vulnerability-transactional stress theory. Psychological Bulletin, 127(6), 773–796.

Hankin, B. L., Abramson, L. Y., Moffitt, T. E., Silva, P. A., McGee, R., & Angell, K. E. (1998). Development of depression from preadolescence to young adulthood: Emerging gender differences in a 10-year longitudinal study. Journal of Abnormal Psychology, 107(1), 128–140. https://doi.org/10.1037/0021-843X.107.1.128

Hare, T. A., Tottenham, N., Galvan, A., Voss, H. U., Glover, G. H., & Casey, B. J. (2008). Biological substrates of emotional reactivity and regulation in adolescence during an emotional go-nogo task. Biological Psychiatry, 63(10), 927–934. https://doi.org/10.1016/j.biopsych.2008.03.015

Hariri, A. R., Mattay, V. S., Tessitore, A., Fera, F., & Weinberger, D. R. (2003). Neocortical modulation of the amygdala response to fearful stimuli. Biological Psychiatry, 53(6), 494–501. https://doi.org/10.1016/S0006-3223(02)01786-9

Hayasaka, S., & Nichols, T. E. (2003). Validating cluster size inference: Random field and permutation methods. NeuroImage, 20(4), 2343–2356.

Heim, C., & Binder, E. B. (2012). Current research trends in early life stress and depression: Review of human studies on sensitive periods, gene–environment interactions, and epigenetics. Experimental Neurology, 233(1), 102–111. https://doi.org/10.1016/j.expneurol.2011.10.032

Herting, M. M., Gautam, P., Spielberg, J. M., Kan, E., Dahl, R. E., & Sowell, E. R. (2014). The role of testosterone and estradiol in brain volume changes across adolescence: A longitudinal structural MRI study: Pubertal Hormones and Brain Volume. Human Brain Mapping, 35(11), 5633–5645. https://doi.org/10.1002/hbm.22575

Hu, S., Pruessner, J. C., Coupé, P., & Collins, D. L. (2013). Volumetric analysis of medial temporal lobe structures in brain development from childhood to adolescence. NeuroImage, 74, 276–287. https://doi.org/10.1016/j.neuroimage.2013.02.032

Hulvershorn, L. A., Cullen, K., & Anand, A. (2011). Toward dysfunctional connectivity: A review of neuroimaging findings in pediatric major depressive disorder. Brain Imaging and Behavior, 5(4), 307–328. https://doi.org/10.1007/s11682-011-9134-3

Jazaieri, H., McGonigal, K., Jinpa, T., Doty, J. R., Gross, J. J., & Goldin, P. R. (2014). A randomized controlled trial of compassion cultivation training: Effects on mindfulness, affect, and emotion regulation. Motivation and Emotion, 38(1), 23–35. https://doi.org/10.1007/s11031-013-9368-z

John, O. P., & Gross, J. J. (2004). Healthy and unhealthy emotion regulation: Personality processes, individual differences, and life span development. Journal of Personality, 72(6), 1301–1333. https://doi.org/10.1111/j.1467-6494.2004.00298.x

Juraska, J. M., & Willing, J. (2017). Pubertal onset as a critical transition for neural development and cognition. Brain Research, 1654, 87–94. https://doi.org/10.1016/j.brainres.2016.04.012

Kaffenberger, T., Brühl, A. B., Baumgartner, T., Jäncke, L., & Herwig, U. (2010). Negative bias of processing ambiguously cued emotional stimuli. Neuroreport, 21(9), 601–605. https://doi.org/10.1097/WNR.0b013e328337ff18

Kessler, R. C., Avenevoli, S., & Ries Merikangas, K. (2001). Mood disorders in children and adolescents: An epidemiologic perspective. Biological Psychiatry, 49(12), 1002–1014. https://doi.org/10.1016/S0006-3223(01)01129-5

Kessler, R. C., Berglund, P., Demler, O., Jin, R., Merikangas, K. R., & Walters, E. E. (2005). Lifetime Prevalence and Age-of-Onset Distributions of DSM-IV Disorders in the National Comorbidity Survey Replication. Archives of General Psychiatry, 62(6), 593–602. https://doi.org/10.1001/archpsyc.62.6.593

Kestenbaum, R. (1992). Feeling happy versus feeling good: The processing of discrete and global categories of emotional expressions by children and adults. Developmental Psychology, 28(6), 1132–1142. https://doi.org/10.1037/0012-1649.28.6.1132

Kim, H., Somerville, L. H., Johnstone, T., Alexander, A. L., & Whalen, P. J. (2003). Inverse amygdala and medial prefrontal cortex responses to surprised faces. NeuroReport, 14(18), 2317.

Kim, H., Somerville, L. H., McLean, A. A., Johnstone, T., Shin, L. M., & Whalen, P. J. (2003). Functional mri responses of the human dorsal amygdala/substantia innominata region to facial expressions of emotion. Annals of the New York Academy of Sciences, 985(1), 533–535. https://doi.org/10.1111/j.1749-6632.2003.tb07120.x

Kim, M. J., Loucks, R. A., Palmer, A. L., Brown, A. C., Solomon, K. M., Marchante, A. N., & Whalen, P. J. (2011). The structural and functional connectivity of the amygdala: From normal emotion to pathological anxiety. Behavioural Brain Research, 223(2), 403–410. https://doi.org/10.1016/j.bbr.2011.04.025

Lang, P. J., Bradley, M. M., & Cuthbert, B. N. (1997). International affective picture system (IAPS): Technical manual and affective ratings. NIMH Center for the Study of Emotion and Attention, 39–58.

Lebron-Milad, K., & Milad, M. R. (2012). Sex differences, gonadal hormones and the fear extinction network: Implications for anxiety disorders. Biology of Mood & Anxiety Disorders, 2(1). https://doi.org/10.1186/2045-5380-2-3

Lee, F. S., Heimer, H., Giedd, J. N., Lein, E. S., estan, N., Weinberger, D. R., & Casey, B. J. (2014). Adolescent mental health—Opportunity and obligation. Science, 346(6209), 547–549. https://doi.org/10.1126/science.1260497

Lewinsohn, P. M., Clarke, G. N., Seeley, J. R., & Rohde, P. (1994). Major Depression in Community Adolescents: Age at Onset, Episode Duration, and Time to Recurrence. Journal of the American Academy of Child & Adolescent Psychiatry, 33(6), 809–818. https://doi.org/10.1097/00004583-199407000-00006

Lupien, S. J., McEwen, B. S., Gunnar, M. R., & Heim, C. (2009). Effects of stress throughout the lifespan on the brain, behaviour and cognition. Nature Reviews Neuroscience, 10(6), 434–445. https://doi.org/10.1038/nrn2639

Mathews, A., Richards, A., & Eysenck, M. (1989). Interpretation of homophones related to threat in anxiety states. Journal of Abnormal Psychology, 98(1), 31–34.

Menzies, L., Goddings, A.-L., Whitaker, K. J., Blakemore, S.-J., & Viner, R. M. (2015). The effects of puberty on white matter development in boys. Developmental Cognitive Neuroscience, 11, 116–128. https://doi.org/10.1016/j.dcn.2014.10.002

Mills, K. L., Goddings, A.-L., Clasen, L. S., Giedd, J. N., & Blakemore, S.-J. (2014). The Developmental Mismatch in Structural Brain Maturation during Adolescence. Developmental Neuroscience, 36(3–4), 147–160. https://doi.org/10.1159/000362328

Monk, C. S., McClure, E. B., Nelson, E. E., Zarahn, E., Bilder, R. M., Leibenluft, E., Charney, D. S., Ernst, M., & Pine, D. S. (2003). Adolescent immaturity in attention-related brain engagement to emotional facial expressions. NeuroImage, 20(1), 420–428. https://doi.org/10.1016/S1053-8119(03)00355-0

Moore, W. E., Pfeifer, J. H., Masten, C. L., Mazziotta, J. C., Iacoboni, M., & Dapretto, M. (2012). Facing puberty: Associations between pubertal development and neural responses to affective facial displays. Social Cognitive and Affective Neuroscience, 7(1), 35–43. https://doi.org/10.1093/scan/nsr066

Muris, P., Meesters, C., & Fijen, P. (2003). The self-perception profile for children: Further evidence for its factor structure, reliability, and validity. Personality and Individual Differences, 35(8), 1791–1802. https://doi.org/10.1016/S0191-8869(03)00004-7

Neta, M., Davis, F. C., & Whalen, P. J. (2011). Valence resolution of facial expressions using an emotional oddball task. Emotion, 11(6), 1425–1433. https://doi.org/10.1037/a0022993

Neta, M., Kelley, W. M., & Whalen, P. J. (2013). Neural responses to ambiguity involve domain-general and domain-specific emotion processing systems. Journal of Cognitive Neuroscience, 25(4), 547–557. https://doi.org/10.1162/jocn_a_00363

Neta, M., Norris, C. J., & Whalen, P. J. (2009). Corrugator Muscle Responses Are Associated With Individual Differences in Positivity-Negativity Bias. Emotion, 9(5), 640–648. https://doi.org/10.1037/a0016819

Neta, M., & Tong, T. T. (2016). Don’t like what you see? Give it time: Longer reaction times associated with increased positive affect. Emotion, 16(5), 730–739. https://doi.org/10.1037/emo0000181

Neta, M., Tong, T. T., & Henley, D. J. (2018). It’s a matter of time (perspectives): Shifting valence responses to emotional ambiguity. Motivation and Emotion, 42(2), 258–266. https://doi.org/10.1007/s11031-018-9665-7

Neta, M., & Whalen, P. J. (2010). The Primacy of Negative Interpretations When Resolving the Valence of Ambiguous Facial Expressions. Psychological Science, 21(7), 901–907. https://doi.org/10.1177/0956797610373934

Ochsner, K. N., Ray, R. D., Cooper, J. C., Robertson, E. R., Chopra, S., Gabrieli, J. D., & Gross, J. J. (2004). For better or for worse: Neural systems supporting the cognitive down-and up-regulation of negative emotion. Neuroimage, 23(2), 483–499. https://doi.org/10.1016/j.neuroimage.2004.06.030

Pagliaccio, D., Luby, J. L., Luking, K. R., Belden, A. C., & Barch, D. M. (2014). Brain-behavior relationships in the experience and regulation of negative emotion in healthy children: Implications for risk for childhood depression. Development and Psychopathology, 26(4), 1289–1303. https://doi.org/10.1017/S0954579414001035

Paus, T., Keshavan, M., & Giedd, J. N. (2008). Why do many psychiatric disorders emerge during adolescence? Nature Reviews Neuroscience, 9(12), 947–957. https://doi.org/10.1038/nrn2513

Perlman, S. B., & Pelphrey, K. A. (2011). Developing connections for affective regulation: Age-related changes in emotional brain connectivity. Journal of Experimental Child Psychology, 108(3), 607–620. https://doi.org/10.1016/j.jecp.2010.08.006

Petersen, A. C., Crockett, L., Richards, M., & Boxer, A. (1988). A self-report measure of pubertal status: Reliability, validity, and initial norms. Journal of Youth and Adolescence, 17(2), 117–133. https://doi.org/10.1007/BF01537962

Petro, N. M., Tong, T. T., Henley, D. J., & Neta, M. (2018). Individual differences in valence bias: FMRI evidence of the initial negativity hypothesis. Social Cognitive and Affective Neuroscience, 13(7), 687–698. https://doi.org/10.1093/scan/nsy049

Pezawas, L., Meyer-Lindenberg, A., Drabant, E. M., Verchinski, B. A., Munoz, K. E., Kolachana, B. S., Egan, M. F., Mattay, V. S., Hariri, A. R., & Weinberger, D. R. (2005). 5-HTTLPR polymorphism impacts human cingulate-amygdala interactions: A genetic susceptibility mechanism for depression. Nature Neuroscience, 8(6), 828–834. https://doi.org/10.1038/nn1463

Pfefferbaum, A., Rohlfing, T., Pohl, K. M., Lane, B., Chu, W., Kwon, D., Nolan Nichols, B., Brown, S. A., Tapert, S. F., Cummins, K., Thompson, W. K., Brumback, T., Meloy, M. J., Jernigan, T. L., Dale, A., Colrain, I. M., Baker, F. C., Prouty, D., De Bellis, M. D., … Sullivan, E. V. (2016). Adolescent Development of Cortical and White Matter Structure in the NCANDA Sample: Role of Sex, Ethnicity, Puberty, and Alcohol Drinking. Cerebral Cortex, 26(10), 4101–4121. https://doi.org/10.1093/cercor/bhv205

Pfeifer, J. H., Kahn, L. E., Merchant, J. S., Peake, S. J., Veroude, K., Masten, C. L., Lieberman, M. D., Mazziotta, J. C., & Dapretto, M. (2013). Longitudinal Change in the Neural Bases of Adolescent Social Self-Evaluations: Effects of Age and Pubertal Development. Journal of Neuroscience, 33(17), 7415–7419. https://doi.org/10.1523/JNEUROSCI.4074-12.2013

Phillips, M. L., Ladouceur, C. D., & Drevets, W. C. (2008). A neural model of voluntary and automatic emotion regulation: Implications for understanding the pathophysiology and neurodevelopment of bipolar disorder. Molecular Psychiatry, 13(9), 833–857. https://doi.org/10.1038/mp.2008.65

Reef, J., Diamantopoulou, S., Meurs, I. van, Verhulst, F. C., & Ende, J. van der. (2011). Developmental trajectories of child to adolescent externalizing behavior and adult DSM-IV disorder: Results of a 24-year longitudinal study. Social Psychiatry and Psychiatric Epidemiology, 46(12), 1233–1241. https://doi.org/10.1007/s00127-010-0297-9

Roy, Amy K., Fudge, J. L., Kelly, C., Perry, J. S. A., Daniele, T., Carlisi, C., Benson, B., Xavier Castellanos, F., Milham, M. P., Pine, D. S., & Ernst, M. (2013). Intrinsic Functional Connectivity of Amygdala-Based Networks in Adolescent Generalized Anxiety Disorder. Journal of the American Academy of Child & Adolescent Psychiatry, 52(3), 290–299.e2. https://doi.org/10.1016/j.jaac.2012.12.010

Roy, Amy Krain, Vasa, R. A., Bruck, M., Mogg, K., Bradley, B. P., Sweeney, M., Bergman, R. L., McClure-Tone, E. B., & Pine, D. S. (2008). Attention bias toward threat in pediatric anxiety disorders. Journal of the American Academy of Child and Adolescent Psychiatry, 47(10), 1189–1196. https://doi.org/10.1097/CHI.0b013e3181825ace

Saarni, C. (1984). An observational study of children’s attempts to monitor their expressive behavior. Child Development, 55(4), 1504–1513. https://doi.org/10.2307/1130020

Samuelsson, H., Jarnvik, K., Henningsson, H., Andersson, J., & Carlbring, P. (2012). The Umeå university database of facial expressions: A validation study. Journal of Medical Internet Research, 14(5). https://doi.org/10.2196/jmir.2196

Schuiling, K. D., & Likis, F. E. (2016). Women’s Gynecologic Health. Jones & Bartlett Learning.

Sewart, A. R., Zbozinek, T. D., Hammen, C., Zinbarg, R. E., Mineka, S., & Craske, M. G. (2019). Positive affect as a buffer between chronic stress and symptom severity of emotional disorders. Clinical Psychological Science. https://doi.org/10.1177/2167702619834576

Silvers, J. A., Insel, C., Powers, A., Franz, P., Helion, C., Martin, R. E., Weber, J., Mischel, W., Casey, B. J., & Ochsner, K. N. (2017). VlPFC–vmPFC–amygdala interactions underlie age-related differences in cognitive regulation of emotion. Cerebral Cortex, 27(7), 3502–3514. https://doi.org/10.1093/cercor/bhw073

Susman, E. J., & Rogol, A. (2013). Puberty and Psychological Development. In R. M. Lerner & L. Steinberg (Eds.), Handbook of Adolescent Psychology (pp. 15–44). John Wiley & Sons, Inc. https://doi.org/10.1002/9780471726746.ch2

Swartz, J. R., Carrasco, M., Wiggins, J. L., Thomason, M. E., & Monk, C. S. (2014). Age-related changes in the structure and function of prefrontal cortex–amygdala circuitry in children and adolescents: A multi-modal imaging approach. NeuroImage, 86, 212–220. https://doi.org/10.1016/j.neuroimage.2013.08.018

Talairach, P., & Tournoux, J. (1988). Co-Planar Stereotaxic Atlas of the Human Brain: 3-Dimensional Proportional System: An Approach to Cerebral Imaging. Thieme Medical.

Tottenham, N., Hare, T. A., Millner, A., Gilhooly, T., Zevin, J., & Casey, B. J. (2011). Elevated amygdala response to faces following early deprivation. Developmental Science, 14(2), 190–204. https://doi.org/10.1111/j.1467-7687.2010.00971.x

Tottenham, Nim, Phuong, J., Flannery, J., Gabard-Durnam, L., & Goff, B. (2013). A negativity bias for ambiguous facial expression valence during childhood: Converging evidence from behavior and facial corrugator muscle responses. Emotion, 13(1), 92–103. https://doi.org/10.1037/a0029431

Tottenham, Nim, Tanaka, J. W., Leon, A. C., McCarry, T., Nurse, M., Hare, T. A., Marcus, D. J., Westerlund, A., Casey, B., & Nelson, C. (2009). The NimStim set of facial expressions: Judgments from untrained research participants. Psychiatry Research, 168(3), 242–249. https://doi.org/10.1016/j.psychres.2008.05.006

Tugade, M. M., & Fredrickson, B. L. (2004). Resilient individuals use positive emotions to bounce back from negative emotional experiences. Journal of Personality and Social Psychology, 86(2), 320–333. https://doi.org/10.1037/0022-3514.86.2.320

van Wingen, G. A., Ossewaarde, L., Bäckström, T., Hermans, E. J., & Fernández, G. (2011). Gonadal hormone regulation of the emotion circuitry in humans. Neuroscience, 191, 38–45. https://doi.org/10.1016/j.neuroscience.2011.04.042

Vantieghem, M. R., Gabard-Durnam, L., Goff, B., Flannery, J., Humphreys, K. L., Telzer, E. H., Caldera, C., Louie, J. Y., Shapiro, M., Bolger, N., & Tottenham, N. (2017). Positive valence bias and parent–child relationship security moderate the association between early institutional caregiving and internalizing symptoms. Development and Psychopathology, 29(2), 519–533. https://doi.org/10.1017/S0954579417000153

Vasey, M. W., Daleiden, E. L., Williams, L. L., & Brown, L. M. (1995). Biased attention in childhood anxiety disorders: A preliminary study. Journal of Abnormal Child Psychology, 23(2), 267–279. https://doi.org/10.1007/BF01447092

Vijayakumar, N., Op de Macks, Z., Shirtcliff, E. A., & Pfeifer, J. H. (2018). Puberty and the human brain: Insights into adolescent development. Neuroscience & Biobehavioral Reviews, 92, 417–436. https://doi.org/10.1016/j.neubiorev.2018.06.004

Vijayakumar, N., Pfeifer, J. H., Flournoy, J. C., Hernandez, L. M., & Dapretto, M. (2019). Affective reactivity during adolescence: Associations with age, puberty and testosterone. Cortex, 117, 336–350. https://doi.org/10.1016/j.cortex.2019.04.024

Wang, F., Kalmar, J. H., He, Y., Jackowski, M., Chepenik, L. G., Edmiston, E. E., Tie, K., Gong, G., Shah, M. P., Jones, M., Uderman, J., Constable, R. T., & Blumberg, H. P. (2009). Functional and structural connectivity between the perigenual anterior cingulate and amygdala in bipolar disorder. Biological Psychiatry, 66(5), 516–521. https://doi.org/10.1016/j.biopsych.2009.03.023

Wechsler, D. (1999). Wechsler Abbreviated Scale of Intelligence. The Psychological Corporation.

Whalen, P. J. (2007). The uncertainty of it all. Trends in Cognitive Sciences, 11(12), 499–500. https://doi.org/10.1016/j.tics.2007.08.016

Whalen, P. J., Davis, F. C., Oler, J. A., Kim, H., Kim, M. J., & Neta, M. (2009). Human amygdala responses to facial expressions of emotion. In The human amygdala. (pp. 265–288). Guilford Press.

Whalen, P. J., Shin, L. M., McInerney, S. C., Fischer, H., Wright, C. I., & Rauch, S. L. (2001). A functional MRI study of human amygdala responses to facial expressions of fear versus anger. Emotion, 1(1), 70–83. https://doi.org/10.1037/1528-3542.1.1.70

Widen, S. C., & Russell, J. A. (2008). Children acquire emotion categories gradually. Cognitive Development, 23(2), 291–312. https://doi.org/10.1016/j.cogdev.2008.01.002

Williams, L. M., Kemp, A. H., Felmingham, K., Liddell, B. J. ., Palmer, D. M., & Bryant, R. A. (2007). Neural biases to convert and overt signals of fear: Dissociation by trait anxiety and depression. Journal of Cognitive Neuroscience, 19(10), 1595–1608. https://doi.org/10.1162/jocn.2007.19.10.1595

Zimmermann, K., Richardson, R., & Baker, K. (2019). Maturational Changes in Prefrontal and Amygdala Circuits in Adolescence: Implications for Understanding Fear Inhibition during a Vulnerable Period of Development. Brain Sciences, 9(3), 65. https://doi.org/10.3390/brainsci9030065

